# LncRNA SNHG16 modulates paclitaxel resistance in breast cancer by regulating Let-7a-5p/DUSP7

**DOI:** 10.1101/2023.08.18.553807

**Authors:** Hui Zhao, Lei Wang, Yueqing Feng, Junzheng Yang

## Abstract

**Objective:** To investigate the effect of long chain non coding RNA (lncRNA) small nucleolar RNA host gene 6 (SNHG16) on paclitaxel (PTX) resistance in breast cancer and to understand its underlying mechanism, to lay a foundation for decreasing the PTX resistance in breast cancer treatment and improving the therapeutic quality for breast cancer patients.

**Methods:** PTX was used to induce the establishment of PTX resistant breast cancer cell lines; the control group (normal cultured MCF-7/PTX cells), si-NC group, si-SNHG16 group, si-SNHG16+anti miR-NC group, and si-SNHG16 +anti-Let-7a-5p group were set to compared the effect of SNHG16 on the PTX resistance in MCF-7 cells; MTT assay, Flow cytometry, and Transwell invasion assay were used to determine the PTX resistance, apoptosis, and invasion ability of MCF-7 cells in different groups, respectively; for further assess the effect of SNHG16 on the PTX resistance, nude mouse tumor transplantation experiment was used; and the potential mechanism of SNHG16 regulated the PTX resistance in MCF-7 cells was explored by double luciferase reporter gene detection method and gene silencing technology.

**Results:** The expression of SNHG16 gene and DUSP7 protein in MCF-7 cell line was the highest, and the expression of Let-7a-5p was the lowest compared with the various breast cancer cell lines (human breast epithelial cell line MCF-10A and human breast cancer cell lines MDA-MB-468 and MDA-MB-453) (P<0.05); PTX could increase the expression of SNHG16 gene and DUSP7 protein, and reduce the expression of Let-7a-5pin MCF-7 cell line (P<0.05); the results of cell experiment and nude mouse transplantation tumor experiment demonstrated that inhibiting the expression of SNHG16 gene could reduce the invasive ability and promote cell apoptosis of MCF-7/PTX cells (P<0.05), and inhibit tumor growth and reduce the PTX resistance in breast cancer transplantation models; simultaneous inhibition of Let-7a-5p and SNHG16 had a weakening effect in MCF-7/PTX cells (P<0.05). Double luciferase reporter gene detection and gene silencing technology demonstrated that inhibiting SNHG16 gene expression could increase the expression of Let-7a-5p and decrease the expression of DUSP7 (P<0.05); inhibiting the Let-7a-5p gene could increase the expression of DUSP7.

**Conclusions:** Inhibiting SNHG16 gene could upregulate Let-7a-5p expression and downregulate the expression of DUSP7 to inhibit MCF-7 cell invasion, promote MCF-7 cell apoptosis, and reduce the PTX resistance in MCF-7 cells and nude mouse tumor transplantation models, those data demonstrated that SNHG16-Let-7a-5p-DUSP7 axis maybe a potential therapeutic strategy for decreasing the PTX resistance in breast cancer in the future.

## 1. Introduction

Breast cancer is one of the common malignant tumor diseases with the high mortality which seriously endangers women’s physical and mental health. Paclitaxel (PTX), as a first-line drug for the treatment of breast cancer, has an obvious effect on improving the anti-tumor activity, but it is prone to drug resistance to affect the treatment, and is becoming one of the big obstacles for the clinical applications of PTX in breast cancer [1]. Therefore, seeking the ideal methods to reduce the PTX resistance of breast cancer cells and understanding the underlying mechanism may help improve the chemotherapy effect and life quality of breast cancer patients.

Many evidences demonstrated that long chain non coding RNA (lncRNA) is involved in the proliferation and metastasis of breast cancer, and participates in drug resistance of breast cancer through various mechanisms [2-3]. Small nucleic RNA host gene 6 (SNHG16) belongs to the lncRNA family and is related to the proliferation, invasion and chemotherapy resistance of many malignant tumor cells including hepatocellular carcinoma [4], pancreatic cancer [5] and colorectal cancer [6]. And the data demonstrated that SNHG16 is highly expressed in breast cancer, and it has been proved to be an important reason for poor prognosis of breast cancer by regulating malignant metastasis, excessive proliferation and chemical resistance in breast cancer cells [7-8]. The accumulated evidences demonstrated that lncRNA including SNHG16 and microRNA could regulate the proliferation, invasion, and chemotherapy sensitivity of cancer cells by binding to miRNA and influencing downstream target gene expression. For example, Shi, et al. demonstrated that downregulation the expression of microRNA Let-7a-5p, a kind of tumor suppressors, could inhibit the proliferation, migration and invasion of triple negative breast cancer cells [9]; it is reported that overexpression of DUSP7 in breast cancer could increase the drug resistance of breast cancer cells, and overexpression of Let-7a-5p could promote the sensitivity of breast cancer cells to PTX by targeting down regulation of DUSP7 [10,11,12]. Whether SNHG16 participates in PTX resistance of breast cancer cell line MCF-7 cells and the potential mechanism relationships among SNHG16, Let-7a-5p and DUSP7 in PTX resistance to breast cancer cell line were not very clear. Therefore, this study aims to explore the effect of SNHG16 on PTX resistance, apoptosis and invasion of in MCF-7 cells and discuss its potential molecular mechanism, hope to provide some ideas to decrease the PTX resistance in breast cancer and improve the therapeutic effect of breast cancer.

## 2. Materials and Methods

### 2.1 Materials and reagents

Human breast epithelial cell line MCF-10A and human breast cancer cell lines MDA-MB-468, MDA-MB-453 and MCF-7 were purchased from Suzhou Beina Chuanglian Biotechnology Company; PTX was purchased from Sigma Company; Penicillin and Streptomycin were purchased from Beijing Leigen Biotechnology Company, DMEM and Fetal bovine serum were purchased from American Biological Industries, MTT detection kit was purchased from Wuhan Huamei Biotechnology Company, Transwell plate was purchased from Corning Company, Lipofectamine^TM^ 2000 kit and reverse transcription kit were purchased from TaKaRa Company, the high purity total RNA rapid extraction kit was purchased from Beijing Biotech Company, the primary antibody DUSP7 and the second antibody coupled with HRP were purchased from the American CST company, the double luciferase Reporter gene kit was purchased from Promega company, and the si-NC (negative control), siRNA SNHG16 (si-SNHG16), SNHG16 overexpression plasmid (pc-SNHG16) and its negative control (pc-NC) were purchased from Shanghai Genepharma company, China. The anti-Let-7a-5p inhibitor (anti-Let-7a-5p) and its negative control (anti-miR-NC), Let-7a-5p mimics and its negative control (miR-NC) were purchased from Ribobio Company.

### 2.2 Methods

#### 2.2.1 Cell culture and establishment of PTX resistant cell lines

Human breast cancer cells were cultured in DMEM medium supplemented with 10% FBS (100U/mL penicillin and 100U/mL streptomycin) in the 37°C, 5% CO _2_ incubator. PTX was dissolved in Dimethyl sulfoxide (DMSO), and MCF-7 cells were treated by PTX to establish the PTX resistant breast cancer cell line.

#### 2.2.2 Cell transfection

MCF-7/PTX cells with the logarithmic growth phase were transfected with Lipofectamine^TM^ 2000 transfection kit. According to the transfection plasmids, the experiment was divided into control group, si-NC group, si-SNHG16 group, si-SNHG16+anti miR-NC group, si-SNHG16+anti Let-7a-5p group, there was six repeats in each group.

#### 2.2.3 qRT-PCR determined the expression of SNHG16, Let-7a-5p, and DUSP7 in MCF-7/PTX cells

The cells from different group were collected after treatment, the total RNA was extracted and reversed into cDNA, then the SNHG16, Let-7a-5p, and DUSP7 gene were amplified by qRT-PCR. The reaction system was as follows: 2μL reverse transcript product, 10μL SYBR Green Mix, 0.5μL forward primers (10μmol/L), and 0.5μL reverse primers (10μmol/L). A total of 45 cycles reaction was amplified as follows: pre-denaturated at 95°C for 5min, denatured at 94°C for 30s, annealing at 60°C for 30s; and used 2^−ΔΔCt^ method to quantify the expression of SNHG16, Let-7a-5p, and DUSP7. The primer sequences of SNHG16, Let-7a-5p, and DUSP7 gene were shown in Table 1.

**Table 1.**
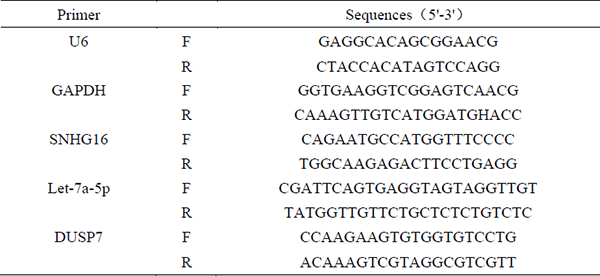
The sequences of primers.

#### 2.2.4 MTT assay

MTT assay was used to detect the apoptosis of MCF-7/PTX cells. Firstly, MCF-7/PTX cells were split into the 96 well plate with the density of 10^4^ cells/mL every well, and cultured for overnight; added different concentrations of PTX (final concentrations are 1, 2, 4, 8, 16, and 32μmol/L), and incubated for 48 hours; added 10μL MTT solution (5 mg/mL) to each well and incubated for 4 hours; finally, added 150μL DMSO to measure the absorbance at a wavelength of 490nm. The cell apoptosis rate is calculated by using the following formula: cell apoptosis rate=[1-A_490 sample_/A_490 control_]×100.

#### 2.2.5 Flow cytometry

MCF-7/PTX cells were collected from each group after treatment and rinsed by cold phosphate buffer (PBS), and the cells from different group were stained by Annexin V, Fluorescein isothiocyanate (FITC), and Propidium iodide (PI) for 15 min; then flow cytometry was used to detect the apoptotic cells.

#### 2.2.6 Transwell invasion test

and MCF-7/PTX cells were split into the Transwell upper chamber (pre-coated with Matrigel) and adjust the concentration of MCF-7/PTX cells to 2×10^4^ per well by using DMEM medium without FBS; added 500μL DMEM medium with 20% FBS to the lower chamber. After incubation at 37°C for 24h, 4% Paraformaldehyde was fixed for 20min, and 1% Crystal violet was dyed for 30min; six visual fields were randomly selected to observe and to count the cell number with a microscope.

#### 2.2.7 Western blot detected the expression of DUSP7 protein

The total protein from each group was extracted according to the specification. and the protein was quantified and separated by SDS-PAGE and transferred to the polyvinylidene fluoride (PVDF) membrane. After sealing the PVDF membrane with skim milk powder for 1 hour, incubated the PVDF membrane in primary antibody DUSP7 and the second antibody coupled with HRP orderly. Finally, the expression of protein was enhanced by chemiluminescence and take photos.

#### 2.2.8 Double luciferase reporter gene detection system verified the target genes

Firstly, Starbase and Targetscan bioinformatics software were used to predict the binding sites of Let-7a-5p and SNHG16 or DUSP7. Two double luciferase reporter vectors, wild-type (WT)-SNHG16 and Mutant (MUT)-SNHG16 were constructed, including the wild-type or mutant binding site of Let-7a-5p. Similarly, constructed WT-DUSP7 3’UTR and MUT-DUSP7 3’UTR dual luciferase reporter vectors with Let-7a-5p binding site fragments. MCF-7/PTX cells were cultured in 96 well plates, and the above-mentioned vectors were transfected into the MCF-7/PTX cells by using Lipofectamine^TM^ 2000; after another 48 hours cultivation, the luciferase activity was measured using a dual luciferase reporter gene detection kit.

In addition, MCF-7/PTX cells were transfected with si-NC, si-SNHG16, pc-NC, and pc-SNHG16 to inhibit or overexpress SNHG16; MCF-7/PTX cells were transfected with anti-Let-7a-5p, anti-miR-NC, Let-7a-5p mimics, and miR-NC to inhibit or upregulate Let-7a-5p expression, and then the mRNA expression of SNHG16, Let-7a-5p, and DUSP7 in MCF-7/PTX cells were detected by qRT-PCR.

#### 2.2.9 Nude mouse tumor transplantation experiment

20 female BALB/c nude mice (4-6 weeks; weighted at 18-25g), were purchased from Beijing Weitong Lihua Experimental Animal Technology Co., Ltd. [license number SCXK (Beijing) 2021-0006], were kept without specific pathogens. 0.1 mL MCF-7/PTX cell suspension (5×10^6^cells/mL) with stable transfection of si-SNHG16 lentivirus plasmid or scramble siRNA was subcutaneously injected into mice, and the mice were divided into four groups: si-NC+PBS group, si-SNHG16+PBS group, si-NC+PTX group, and si-SNHG16+PTX group. After subcutaneous transplantation for 8 days, nude mice were injected with 0.1mL PBS or 0.1mL paclitaxel (3mg/kg, tail vein injection, dissolved in 0.1mL of PBS) [13], administered every 3 days for 30 days, monitored the growth of xenograft tumor of breast cancer in nude mice every 7 days, and calculated the tumor volume according to the formula: (length×width^2^)/2. Nude mice were euthanized, umor tissue was removed and weighed at 30^th^ day, and the tumor tissue was stored at -80°C until for use.

### 2.3 Statistical Analysis

The measurement data were expressed as mean±standard deviation (SD), and SPSS 20.0 software was used for statistical analysis. t-test was used for comparison between the two groups, and one-way ANOVA was used for comparison among multiple groups. P<0.05 was considered statistically differences.

## 3 Results

### 3.1 Compared the expression of SNHG16, Let-7a-5p, and DUSP7 in various cell lines

The results of qPCR and western blotting demonstrated that the expression of SNHG16 and DUSP7 in MCF-7 cell line was the highest, and the expression of Let-7a-5p were the lowest (P<0.05) compared with MCF-10A cell line, MDA-MB-468 cell line, and MDA-MB-453 cell line. After PTX treatment, the expressions of SNHG16 and DUSP7 in MCF-7/PTX cells were higher, while the expression of Let-7a-5p were lower than those in MCF-7 cells (P<0.05) (Table 2).

**Table 2.**
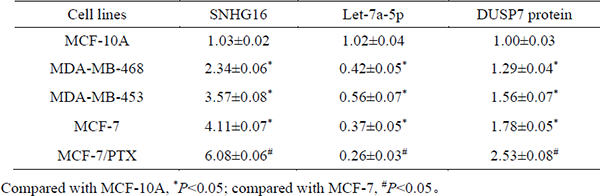
The comparison of expression of SNHG16?Let-7a-5p ? DUSP7 in different cell lines(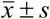, n=6)

| Cell lines | SNHG16 | Let-7a-5p | DUSP7 protein |
| --- | --- | --- | --- |
| MCF-10A | 1.03±0.02 | 1.02±0.04 | 1.00±0.03 |
| MDA-MB-468 | 2.34±0.06* | 0.42±0.05* | 1.29±0.04* |
| MDA-MB-453 | 3.57±0.08* | 0.56±0.07* | 1.56±0.07* |
| MCF-7 | 4.11±0.07* | 0.37±0.05* | 1.78±0.05* |
| MCF-7/PTX | 6.08±0.06<sup>#</sup> | 0.26±0.03<sup>#</sup> | 2.53±0.08<sup>#</sup> |

Compared with MCF-10A, \* $P<0.05$ ; compared with MCF-7, <sup>#</sup> $P<0.05$ 。

**Figure 1.**
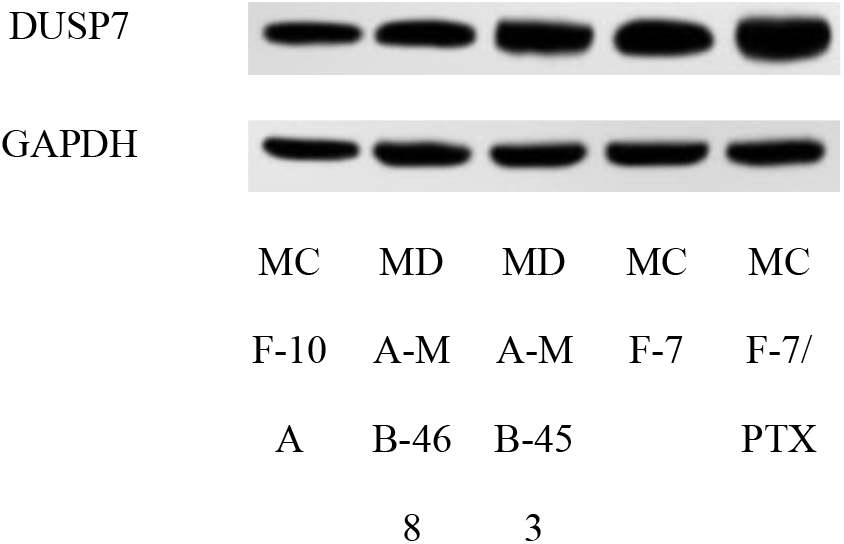
Western blot detected the protein expression of DUSP7 in different cell lines.

### 3.2 Inhibiting the expression of SNHG16 could reduce the invasion and proliferation of MCF-7 cells, simultaneously inhibiting SNHG16 gene and Let-7a-5p gene has an antagonistic effect

The results demonstrated that the expression of SNHG16 was decreased and the expression of Let-7a-5p was increased, and invasive capability of MCF-7 cells were decreased and the cell apoptosis rate of MCF-7 cells were significantly increased in si-SNHG16 group, si-SNHG16+anti-miR-NC group and si-SNHG16+anti Let-7a-5p group compared with the control group and si-NC group (P<0.05); compared with si-SNHG16 group and si-SNHG16+anti-miR-NC group, the expression of Let-7a-5p and apoptosis rate of MCF-7 cells were decreased, while invasive capability were significantly increased (P<0.05) in the si-SNHG16+anti Let-7a-5p group, those data demonstrated that inhibition of SNHG16 gene could decrease the invasion and proliferation of MCF-7 cells, but there was an antagonistic effect between SNHG16 gene and Let-7a-5p gene (Figure 2, Figure 3, and Table 3).

**Figure 3.**
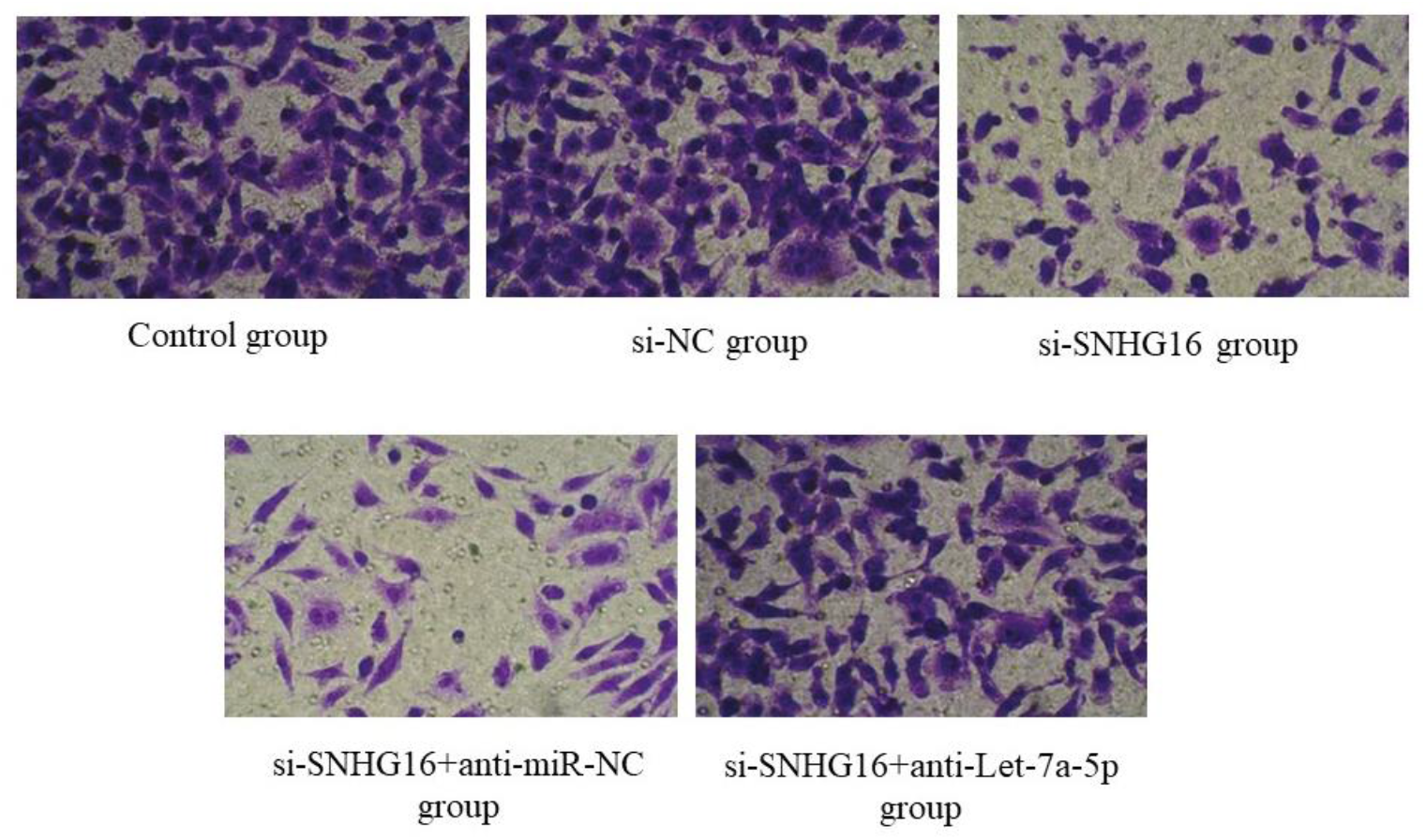
Transwell experiment detected the cell invasion of MCF-7 cells in each group.

**Figure 4.**
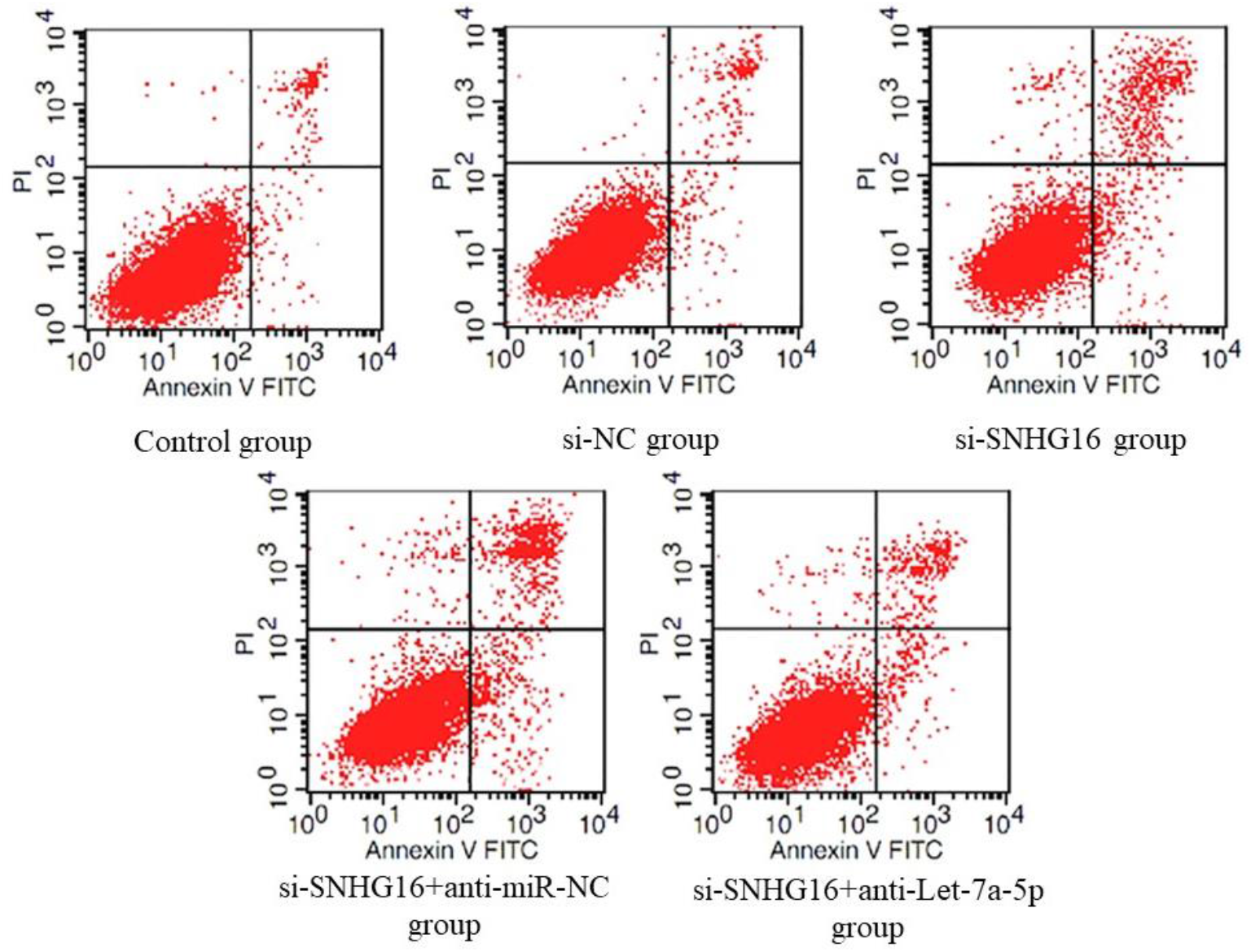
Flow cytometry detected the cell apoptosis of MCF-7 cells in each group.

**Table 3.**
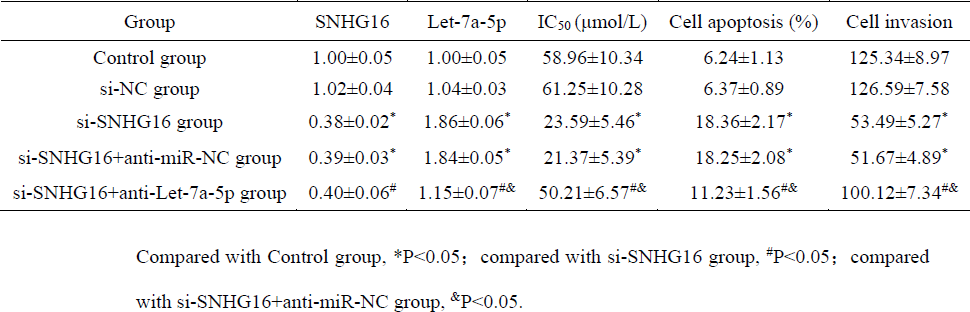
Comparison of the expression of SNHG16, Let-7a-5p, IC50, cell apoptosis and invasion capability of MCF-7 cells in different group.

### 3.3 Silencing the expression of SNHG16 gene in MCF-7 cells could downregulate the expression of Let-7a-5p and upregulate the DUSP7 to inhibit tumor growth and decrease the PTX-induced drug resistance in breast cancer transplantation model

The data demonstrated that the tumor volume in different group was increased with the time; compared with si-NC+PBS group, the tumor volume and tumor weight in si-NC+PTX group, si-SNHG16+PBS group and si-SNHG16+PTX group were decreased significantly (P<0.05); compared with si-NC+PBS group, si-NC+PTX group, and si-SNHG16+PBS group, the tumor volume and tumor weight in si-SNHG16+PTX group were decreased significantly (P<0.05).

**Table 4.** Comparison of the tumor volume and tumor weight in breast cancer transplantation model in each group (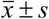,n =5)

| Group | Tumor volume (mm<sup>3</sup>) |  |  |  | Tumor weight (g) |
| --- | --- | --- | --- | --- | --- |
|  | 7d | 14d | 21d | 28d |  |
| si-NC+PBS | 51.49±7.22 | 171.38±26.64 | 264.03±35.22 | 395.10±47.38 | 0.68±0.07 |
| si-SNHG16+PBS | 46.27±6.03 | 109.25±18.70* | 150.45±24.15* | 187.42±30.21* | 0.39±0.05* |
| si-NC+PTX | 49.86±6.79 | 134.06±21.35* | 207.30±27.28* | 296.60±38.57* | 0.54±0.06* |
| si-SNHG16+PTX | 45.70±6.31 | 72.12±13.09<sup>#&</sup> | 96.74±18.12<sup>#&</sup> | 125.13±20.30<sup>#&</sup> | 0.26±0.04<sup>#&</sup> |

Compared with si-NC+PBS group, \*P<0.05; compared with si-SNHG16+PBS group, <sup>#</sup>P<0.05; compared with si-NC+PTX group, <sup>&</sup>P<0.05.

**Table 5.**
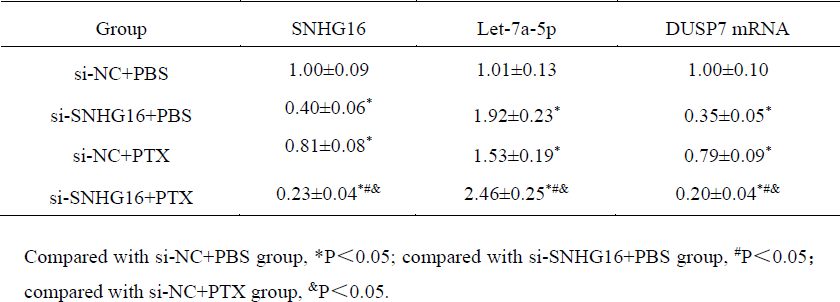
Comparison of the expression of SNHG16, Let-7a-5p and DUSP7 in breast cancer transplantation model in each group(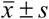, n=5)

| Group | SNHG16 | Let-7a-5p | DUSP7 mRNA |
| --- | --- | --- | --- |
| si-NC+PBS | 1.00±0.09 | 1.01±0.13 | 1.00±0.10 |
| si-SNHG16+PBS | 0.40±0.06* | 1.92±0.23* | 0.35±0.05* |
| si-NC+PTX | 0.81±0.08* | 1.53±0.19* | 0.79±0.09* |
| si-SNHG16+PTX | 0.23±0.04*#& | 2.46±0.25*#& | 0.20±0.04*#& |

Compared with si-NC+PBS group, \*P<0.05; compared with si-SNHG16+PBS group, #P<0.05; compared with si-NC+PTX group, &P<0.05.

### 3.4 SNHG16 could regulate the Let-7a-5p and DUSP7 to decrease the invasion and proliferation of MCF-7 cells and decrease the PTX-induced drug resistance of MCF-7 cells

Starbase software and Targetscan software predicted that there were binding sites between Let-7a-5p and SNHG16, while there were binding sites between DUSP7 and Let-7a-5p (Figure 5). The results of double luciferase reporter gene detection demonstrated that the luciferase activity in the Let-7a-5p mimics+SNHG16-WT group was significantly decreased (P<0.05), while there was no statistically significant difference in luciferase activity between the miR-NC+SNHG16-MUT group and the Let-7a-5p mimics+SNHG16-MUT group (P>0.05) compared with the miR-NC+SNHG16-WT group; Compared with the miR-NC+DUSP7 3’UTR-WT group, the Let-7a-5p mimics+DUSP7 3’UTR-WT group demonstrated a significant decrease in luciferase activity (P<0.05), while there was no statistically significant difference in luciferase activity between the miR-NC+DUSP7 3’UTR-MUT group and the Let-7a-5p mimics+DUSP7 3’UTR-MUT group (P>0.05).

Compared with the si-NC group, the expression of SNHG16 in the si-SNHG16 group was decreased, while the expression of Let-7a-5p was increased (P<0.05); compared with the pc-NC group, the expression level of SNHG16 in the pc-SNHG16 group increased, while the expression level of Let-7a-5p decreased (P<0.05); Compared with the anti miR-NC group, the expression of Let-7a-5p were decreased and the expression of DUSP7 was increased in anti-Let-7a-5p group (P<0.05); compared with the miR-NC group, the expression Let-7a-5p was increased and expression DUSP7 was decreased in the Let-7a-5p mimics group (P<0.05).

**Figure 5.**
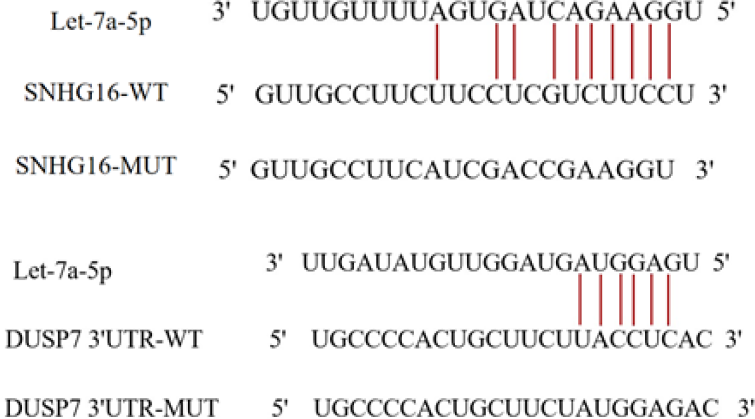
The prediction result of binding sites between Let-7a-5p and SNHG16, and binding sites between Let-7a-5p and DUSP7.

**Table 5.** Targeting relationship between Let-7a-5p and SNHG16 detected by double luciferase Reporter gene (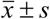, n=6)

| Group | luciferase activity |
| --- | --- |
| miR-NC+SNHG16-WT | 1.00±0.08 |
| miR-NC+SNHG16-MUT | 0.99±0.07 |
| Let-7a-5p mimics+SNHG16-WT | 0.34±0.05* |
| Let-7a-5p mimics+SNHG16-MUT | 1.01±0.06 |

Compared with miR-NC+SNHG16-WT group, \* $P<0.05$

**Table 6.** Targeting relationship between Let-7a-5p and DUSP7 detected by double luciferase Reporter gene (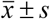, n=6)

| Group | luciferase activity |
| --- | --- |
| miR-NC+DUSP7 3'UTR-WT | 1.00±0.05 |
| miR-NC+DUSP7 3'UTR-MUT | 0.98±0.06 |
| Let-7a-5p mimics+DUSP7 3'UTR-WT | 0.35±0.04* |
| Let-7a-5p mimics+DUSP7 3'UTR-MUT | 1.01±0.05 |

Compared with miR-NC+DUSP7 3'UTR-WT group, \* $P<0.05$ .

**Table 7.** SNHG16 regulate the expression of Let-7a-5p in different group (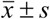, n=6)

| Group | SNHG16 | Let-7a-5p |
| --- | --- | --- |
| si-NC | 1.00±0.09 | 1.01±0.10 |
| si-SNHG16 | 0.36±0.05* | 1.92±0.14* |
| pc-NC | 1.02±0.13 | 0.99±0.08 |
| pc-SNHG16 | 2.89±0.20<sup>#</sup> | 0.40±0.05<sup>#</sup> |

Compared with si-NC group, \* $P<0.05$ ; Compared with pc-NC group, <sup>#</sup> $P<0.05$ .

**Table 8.** Let-7a-5p regulate the expression of DUSP7 in different group (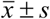, n=6)

| Group | DUSP7 |
| --- | --- |
| Control | 1.00±0.11 |
| si-NC | 1.01±0.12 |
| si-SNHG16 | 0.37±0.05* |
| si-SNHG16+anti-miR-NC | 0.36±0.05 |
| si-SNHG16+anti-Let-7a-5p | 0.74±0.09<sup>#&</sup> |

Compared with Control, \* $P<0.05$ ; compared with si-SNHG16 group, <sup>#</sup> $P<0.05$ ; compared with si-SNHG16+anti-miR-NC group, <sup>&</sup> $P<0.05$ .

## 4 Discussion

Breast cancer is caused by abnormal proliferation of breast epithelial cells, breast cancer cells could be transferred through body fluid, which result in the high fatality rate of breast cancer. It is reported that PTX is a kind of natural anti-cancer drugs for ovarian cancer, breast cancer, and bladder cancer, but it is easy to develop into drug resistance, which seriously hinders clinical application of PTX [14]. The evidences demonstrated that lncRNA including SNHG16 played the crucial role in the chemotherapy resistance of various tumors [15-16], SNHG16 was located on human chromosome 17, and was considered as a potential carcinogen of breast cancer [17]. In our data, we found that the expression of SNHG16 in MCF-7 cells was the highest, and silencing SNHG16 in MCF-7 cells could increase the sensitivity of MCF-7/PTX cells to PTX, promote cell apoptosis, and inhibit the invasion of MCF-7 cells, and our data demonstrated that SNHG16 participated in the process of carcinogenesis and development of breast cancer, and inhibiting the expression of SNHG16 could play an important role in MCF-7 cells and may become a potential treatment target for breast cancer.

Let-7a-5p is a kind of genes which was widely studied and it was reported that Let-7a-5p could inhibit cancer cell proliferation and promote apoptosis [18-19]. The evidences demonstrated that the expression of Let-7a-5p in breast cancer tissues was decreased, and Let-7a-5p could downregulate expression of CC Chemokine receptor 7 (CCR7) to inhibit the migration and invasion of breast cancer cells [19], and Yang et al. [12] demonstrated that the expression of Let-7a-5p in PTX resistant breast cancer tissues and breast cancer cells was decreased than that in PTX sensitive breast cancer tissues and parental breast cancer cells. Those evidences demonstrated that Let-7a-5p may play an important role in process of cancer cell proliferation and invasion capability. In our data, the results of double luciferase reporter gene detection and qPCR demonstrated SNHG16 could regulate expression of Let-7a-5p, silencing SNHG16 could increase the expression of Let-7a-5p and improve the sensitivity of breast cancer cells to PTX; and inhibiting the expression of Let-7a-5p could eliminate the effect of SNHG16 silencing on PTX sensitivity and apoptosis, and inhibiting the invasion of PTX resistant breast cancer cells. those data suggested that silencing SNHG16 may increase the sensitivity of breast cancer cells to PTX, induce apoptosis and inhibit cell invasion by upregulating the expression of Let-7a-5p.

DUSP7 is a member of the DUSP family, which is highly expressed in breast cancer and plays the role of proliferation and invasion of cancer cells [20], and the evidences demonstrated that high expression of Let-7a-5p could enhance the sensitivity of breast cancer cells to PTX by downregulating the expression of DUSP7 [12]. Our data also demonstrated the same results in MCF-7 cells, and the results of breast cancer transplantation model demonstrated that silencing the SNHG16 could upregulate the expression of Let-7a-5p and downregulate DUSP7 to affect the progression of breast cancer. Those evidences indicated that SNHG16 could affect the process of carcinogenesis of breast cancer by regulating the expression of Let-7a-5p and DUSP7.

## 5 Conclusion

Silencing SNHG16 could upregulate Let-7a-5p expression and downregulate DUSP7 expression to increase the sensitivity of MCF-7 cells to PTX, inhibit MCF-7 cells invasion, increase MCF-7 cells apoptosis, inhibit tumor growth and decrease the PTX-induced drug resistance in breast cancer transplantation model. Those data may provide new insights for developing an effective strategy to alleviate PTX resistance of breast cancer patients in the future.

## 6 Ethical declarations

The animal experiments were approved by ethics committee of The First Affiliated Hospital of Xinxiang Medical University & the First Clinical College of Xinxiang Medical University.

## 7 Human and animal rights

The study is according to the Guide for the Care and Use of Laboratory Animals.

## 8 Acknowledgements

None.

